# Targeted knockdown of ribulose-1, 5-bisphosphate carboxylase-oxygenase in rice mesophyll cells impact on photosynthesis and growth

**DOI:** 10.1101/2020.08.20.259382

**Authors:** Chirag Maheshwari, Robert A Coe, Shanta Karki, Sarah Covshoff, Ronald Tapia, Aruna Tyagi, Julian M. Hibberd, Robert T. Furbank, W Paul Quick, Hsiang-Chun Lin

**Affiliations:** C_4_ Rice Centre, International Rice Research Institute (IRRI), Los Baños, Philippines; Department of Plant Sciences, University of Cambridge, Cambridge, CB2 3EA, United Kingdom; Division of Biochemistry, ICAR-Indian Agricultural Research Institute, New Delhi, India; ARC Centre of Excellence for Translational Photosynthesis, Research School of Biology, The Australian National University, Acton, 2601, Australia; Department of Animal and Plant Sciences, University of Sheffield, Sheffield, S10 2TN, UK

**Keywords:** *Oryza sativa*, antisense, Rubisco, RBCS, C_4_ rice

## Abstract

We generated antisense constructs targeting two of the five Rubisco small subunit genes (*OsRBCS2 and 4*) which account for between 30-40% of the RBCS transcript abundance in leaf blades. The constructs were driven by a maize phosphoenolpyruvate carboxylase (PEPC) promoter known to have enriched expression in mesophyll cells (MCs). In the resulting lines leaf Rubisco protein content was reduced by between 30-50% and CO_2_ assimilation rate was limited under photorespiratory and non-photorespiratory conditions. A relationship between Rubisco protein content and CO_2_ assimilation rate was found. This was associated with a significant reduction in dry biomass accumulation and grain yield of between 37 to 70%. In addition to serving as a resource for reducing Rubisco accumulation in a cell-preferential manner, these lines allow us to characterize gene function and isoform specific suppression on photosynthesis and growth. Our results suggest that the knockdown of multiple genes is required to completely reduce Rubisco accumulation in MCs.

## Introduction

In C_3_ photosynthesis the first step of CO_2_ fixation is carried out by ribulose-1, 5-bisphosphate carboxylase-oxygenase (Rubisco, EC 4.1.1.39) in the chloroplast of mesophyll cells (MCs). Photosynthetic rate is limited by the activity of Rubisco because of its extremely low catalytic turnover rate and competing oxygenation reaction, which leads to the formation of toxic metabolites which must be broken-down by a series of reactions in a process known as photorespiration (von Caemmerer and Quick, 2000). This consumes energy leading to the loss of previously fixed CO_2_. As the temperature increases the rate of oxygenation increases due to a decline in the ratio of CO_2_ to O_2_ in the leaf cell, as such rates of photorespiration are highest in hot climates such as the major rice growing regions of the world. To compensate for these catalytic inefficiencies, Rubisco is produced in large amounts accounting for up to 50% of total leaf protein and forming a major sink for leaf nitrogen (Parry et al., 2013).

Some species suppress the oxygenation reaction allowing Rubisco to operate close to its maximal carboxylase rate (Whitney et al., 2011). In C_4_ plants Rubisco is localized to the bundle sheath cells (BSCs) allowing CO_2_ to be concentrated at high levels at the exclusion of oxygen. As a result, radiation use efficiency (RUE) and yield of C_4_ species such as maize and sorghum can be up to 50% higher than those of C_3_ species such as rice and wheat (Sheehy et al., 2007). Water use efficiency (WUE) and nitrogen use can be up to 50% lower (Ghannoum et al., 2011). The potential efficiencies associated with C_4_ photosynthesis has stimulated interest in converting rice (*Oryza sativa* L.) from a C_3_ to a C_4_ photosynthetic pathway (Hibberd et al., 2008, Kajala et al., 2011; Ermakova et al. 2019). The C_4_ Rice Consortium (https://c4rice.com/; von Caemmerer et al., 2012) was established with this goal in mind. To investigate the feasibility of such an endeavor a toolkit of transgenic resources is being assembled (Kajala et al., 2011; Ermakova et al., 2019). Among these, there are lines in which parts of the photorespiratory cycle have been selectively downregulated to test the hypothesis of whether this primes a plant for C_4_ photosynthesis (Lin et al., 2016). Here we report on the downregulation and translocation of Rubisco from MCs to BSCs and the effect on rice photosynthesis and growth.

In rice the Rubsico holoenzyme is made up of eight large Rubisco subunits (rbcLs) encoded by a single gene on the chloroplast DNA and eight small Rubisco subunits (RBCSs) encoded by a five multigene family on the nucleus DNA (Dean et al., 1989; Rodermel 1999; Sasanuma 2001). *OsRBCS1* is located on chromosome 2 but is not expressed in the leaf blade (Morita et al., 2014; Suzuki et al., 2007, 2009). The remaining four genes (*OsRBCS2-OsRBCS5*) are in a tandem array on chromosome 12 and are highly expressed in photosynthetic active tissues such as leaf blades (Suzuki et al., 2009). The deduced amino acid sequence without the transit peptide for targeting the chloroplast are completely identical, leading to the assumption that there are no functional differences within the *RBCS* gene family within a species. It has previously been shown that suppression of a *RBCS* gene leads to a reduction in the holoenzyme (Hudson et al., 1992; Makino et al., 1997; Rodermel et al., 1988) with the expression of *rbcL* regulated at its transcript level in response to the availably of RBCS protein (Makino et al., 1997; Suzuki and Makino 2012). Although RBCS protein has no catalytical function for CO_2_ fixation, it is important for maximal activity and structural stability for the Rubisco holoenzyme (Andersson and Backlund, 2008).

To reduce Rubisco protein accumulation, we generated antisense constructs targeting two of the five *RBCS* genes (*OsRBCS2* and *4*) expressed them in photosynthetic tissue. Together these account for between 30-40% of the *RBCS* transcript abundance in leaf blades (Suzuki et al., 2009), allowing a substantial reduction in Rubisco protein accumulation to be achieved without a lethal phenotype developing. Lines must produce viable seed in order to be crossed with lines expressing C_4_ enzyme genes facilitating gradual replacement of C_3_ with C_4_ photosynthesis (Kajala et al., 2011). The antisense genes against *OsRBCS2* and *4* are driven by a maize phosphoenolpyruvate carboxylase (PEPC) promoter known to have enriched expression in MCs (Matsuoka et al., 1994). In addition to serving as a resource for reducing Rubisco accumulation in a cell-specific manner, these lines allow us to characterize gene function and isoform specific suppression on photosynthesis and growth.

## Materials and Methods

### Generation of *OsRBCS2* and *OsRBCS4* knockdown transgenic rice lines

To individually reduce *OsRBCS2* (Os12g17600) and *OsRBCS4* (Os12g19470) expression in rice MCs, the full coding sequences (CDS) of LOC_Os12g17600.1 and LOC_Os12g19470.2 were cloned separately in the antisense direction via RT-PCR using the following primers: 5’– CACCTTAGTTGCCACCAGACTCCT and 5’–ATGGCCCCCTCCGTGATGGC for *OsRBCS2*, 5’–CACCTTAGTTGCCGCCTGACTCCT and 5’–ATGGCTCCCTCGGTGATGGC for *OsRBCS4*. The CACC at the 5’ end of the forward primer allowed directional cloning into pENTR/D-TOPO (ThermoFisher Scientific, USA) and subsequent Gateway cloning into the pSC110 expression vector. The pSC110 vector was created by Gibson Assembly and contained the B73 *ZmPEPC* promoter to drive gene expression in rice MCs (Gibson et al., 2009; Osborn et al., 2017; Matsuoka et al., 1994). Both antisense *OsRBCS2* and *OsRBCS4* vectors were verified by sequencing and transformed into rice (*Oryza sativa* indica *cv*. IR64) using *Agrobacterium*-mediated transformation following the method described by Yin et al. (2019).

### Plant growth conditions

Plants were grown at the International Rice Research Institute (IRRI), Los Baños, Philippines, 14°9’53.58”N, 121°15’32.19”E. Seeds were sown and germinated in 100 ml rootrainers (http://rootrainers.co.uk/), after one-week plants were transplanted into 7-liter pots filled with soil from the IRRI upland farm. Plants were grown in a containment transgenic screen house with a day/night temperature of 35 ± 3°C/28 ± 3°C and relative humidity of 70-90%. Maximum light intensity was 2000 µmol photons m^−2^ s^−1^ on a clear sunny day.

### PCR screening

Transgenic plants were subjected to genomic PCR screening to confirm the presence of *OsRBCS* antisense DNA sequence. PCR was carried out by using the KAPA 3G plant PCR kit (Kapa Biosystem, USA; https://www.sigmaaldrich.com/life-science/roche-biochemical-reagents/kapa-genomics-reagents.html) with the gene-specific primers (5’-ACGACTCCCCATCCCTATTT and 5’-TGCATTCGTCCGTATCATC for *OsRBCS2*; 5’-CATTGATCACCAATCGCATC and 5’-ACGTCAGCAATGGCGGAA for *OsRBCS4*). Plasmid DNA containing antisense *OsRBCS2* and *OsRBC4* were used as positive controls and non-transgenic rice and water as negative controls. PCR conditions were as follows: pre-denaturation for 5 min at 95°C, 30 cycles of the polymerization reaction consisting of a denaturation step for 20 s at 95°C, an annealing step for 15 s at 60°C and an extension step for 1 min at 72 °C, and a final extension step for 5 min at 72°C.

### DNA blot analysis

Large-scale genomic DNA was extracted from leaves at the mid-tillering stage. DNA blot analysis was carried out as described by Lin et al., (2016) except genomic DNA was digested with *BgL*II restriction endonuclease (New England Biolabs, USA; https://international.neb.com/).

### Quantitative RT-PCR

Total RNA was extracted and analyzed as described in Lin et al., (2016) except for the primer pairs used. The mRNA level of each *OsRBCS* gene was quantified on a fold change basis in comparison with wild-type plants. 3’-untranslated regions (UTRs) of the *OsRBCS* gene family have different nucleotide sequences and so were utilized for amplification. The *Elongation factor 1-alpha* gene (*OsEF1α*; Os03g0177500) was used as an internal standard. Primers pairs were: 5’-CTTCGGCAACGTCAGCAATG and 5’-AGCACGGCCGGTAAAATCA for *OsRBCS2*; 5’-TTCCAAGGGCTCAAGTCCAC and 5’-ATAGGCAGCAATCCACCAGC for *OsRBCS3*; 5’-GTGGCCGATTGAGGGCATTA and 5’-TCACCGTCACCGAACGATAG for *OsRBCS4*; 5’-GGTGTGGCCGATTGAAGG and 5’-ACTGGGAAAGGGGAAACCAAT for *OsRBCS5*; 5’-CCACTGGTGGTTTTGAGGC and 5’GGCCCTTGTACCAGTCAAG for *OsEF1α*.

### Soluble leaf protein

Leaf samples for soluble protein were harvested between 09:00 and 11:00 h from the fourth fully expanded leaf. Proteins were extracted and fractionated as described previously Lin et al. (2016). Samples were loaded based on an equal leaf area (0.396 mm^2^ for rbcL and RBCS). After electrophoresis, proteins were electroblotted onto a polyvinylidene difluoride membrane and probed with antisera against Rubisco protein (provided by Richard Leegood, Sheffield University, UK) at a dilution 1:200. A peroxidase-conjugated secondary antibody was used at a dilution of 1:200 and immunoreactive bands were visualized with ECL Western Blotting Detection Reagents (GE Healthcare, USA; https://www.gelifesciences.com). Rubisco protein content was quantified from SDS-page gels using ALPHA-Ease FC software (Alpha Innotech, USA), values are expressed as the percentage protein reduction compared to the wild-type.

### Leaf chlorophyll content, plant growth analysis and destructive harvesting

Leaf chlorophyll content was measured with a SPAD 502 Chlorophyll Meter (SPAD, Konica Minolta; https://www.konicaminolta.com) at the mid-tillering stage on the upper fully expanded leaves. Values given are the average ± SE of three measurements from six plants per line. Plant height was measured from the soil surface to the tip of the youngest fully expanded leaf. Total tiller number was counted prior to harvesting for destructive measurements. All above-ground biomass (leaves, stems, and sheaths) were harvested, weighed, and placed in paper bags, and oven dried at 70°C until a constant dry biomass weight was achieved. Values of plant height, total tiller number, and dry biomass are presented as the average ± SE of ten plants per line.

### Gas exchange measurement

Leaf gas exchange measurements were made at IRRI using a LI-6400XT portable photosynthesis system (LI-COR Biosciences, USA; https://www.licor.com) as described in Lin et al. (2016) during the tillering stage for each line and wild type, 60-65 days post-germination.

### Immunolocalization

Leaf samples were harvested between 09:00 h and 11:00 h from 9-week-old plants. The middle portion of the seventh fully expanded leaf was dissected and fixed for four hours in a solution containing 4% paraformaldehyde, 0.2% glutaraldehyde, and 25 mM sodium phosphate buffer pH 7.2. After fixation, sections were rinsed four times in 25 mM phosphate buffer over a period of 60 min. Thin leaf sections were cut using a razor blade and blocked in TBST buffer (0.1% Tween 20, 20mM Tris, 154 mM NaCl) containing 3% milk for two hours at room temperature. Sections were probed with antisera against Rubisco protein diluted 1:100 in blocking solution and incubated overnight at 4°C. The remaining steps were performed at room temperature. Sections were washed six times with blocking solution and then incubated for two hours with Alexa Fluor 488 goat anti-rabbit IgG (Invitrogen, USA; https://www.thermofisher.com/ph/en/home/brands/invitrogen.html) at 37°C in the dark. Sections were washed six times with blocking solution, post-stained with 0.05% calcofluor white for five min and washed with distilled water. Sections were mounted on microscope slides in 50% glycerol and examined on a BX61 Disk Scanning microscope (Olympus, USA; https://www.olympus-global.com) with florescence function under DAPI, RFP and GFP filters.

### Leaf Anatomy Analysis

Leaf width was measured at the middle portion of the fully expanded penultimate leaf. Between 2 to 4 mm^2^ of the middle portion of the penultimate leaf was cut and fixed in formaldehyde-acetic acid-alcohol (FAA) solution under vacuum (20 psi) at room temperature for at least 12 h. Leaf sections were rinsed twice in water for 60 min. Leaf sections were dehydrated using an ethanol series, incubated twice in 70% ethanol for 30 min; once in 80% ethanol for 30 min; once in 90% ethanol for 30 min, and finally three times in 100% ethanol for 30 min. Leaf sections were infiltrated in 10% ethanol and varying concentrations of Spurr’s resin solution (10% to 100%). Sections were incubated for at least 60 min at each concentration solution. Leaf sections were then finally infiltrated twice for 4 h with 100% Spurr’s resin solution. Sections were placed in fresh 100% Spurr’s resin solution in molds. Samples were polymerized overnight at 70°C. Embedded leaf sections were cut into 10-15 μm thick sections on a microtome (MT2-B, DuPont-Instruments-Sorvall, USA) and then dried. Sections were stained with 0.05% toluidine blue O stain in 0.1% sodium carbonate, pH 11.1 for 5 min then rinsed four times with distilled water, each for 5 min. Sections were dried and mounted on slides using Permount^TM^ (Fischer Scientific, USA; https://www.fishersci.com/us/en/home.html). Transverse images were acquired using a BX51 Olympus microscope (Olympus, USA; https://www.olympus-global.com/) at 4X and 40X magnification. Mesophyll cell number was counted by calculating the numbers of mesophyll cells in between minor veins. Leaf thickness, interveinal distance (IVD), vein number (major and minor), mesophyll cell length was measured as described in Chatterjee et al. (2016).

### Statistics

Statistical analyses were performed using Statistical Tool for Agricultural Research (STAR) software (International Rice Research Institute, Philippines) using a one-way analysis of variance (ANOVA) or a Student’s t-test with a P-value of < 0.05.

## Results

### *OsRBCS2* and Os*RBCS4* specific knockdown lines

A total of 190 T_0_ plants were PCR positive for the antisense-Os*RBCS2* construct and 30 T_0_ plants for the antisense-*OsRBCS4* construct. Plants with reduced Rubisco protein accumulation relative to the wild-type rice were selected for DNA blot analysis in order to confirm T-DNA integration. Two *OsRBCS2* knockdown events (*rbcs2*-165 and *rbcs2*-266) and two *OsRBCS4* knockdown events (*rbcs4*-022 and *rbcs4*-053) with the lowest Rubisco accumulation were selected from T_1_ progeny. These were advanced generating a total of 40 PCR positive T_3_ generation *OsRBCS2* knockdown plants (from two events *rbcs2-*165 and *rbcs2*-266) and 40 T_2_ generation *OsRBCS4* knockdown plants (from two events *rbcs4*-022 and *rbcs4*-053) (Fig. S1). DNA blot analysis showed that these transgenic lines carried between two and four copies of the antisense constructs (Fig. S2). This led to a reduction in the accumulation of *OsRBCS2* transcripts of between 15 and 24-fold (Fig. 1A) and *OsRBCS4* transcripts of between 6 and 12-fold in the respective knockdown lines (Fig. 1C). Reduction of either isoform was also associated with a slight increase in the accumulation of *OsRBCS3* and *OsRBCS5* transcripts (Fig 1B & 1D). These results indicate that the *OsRBCS2* and *OsRBCS4* genes were selectively suppressed by the antisense approach. Correspondingly immunoblot analysis showed a reduction in the accumulation of RBCS in both *OsRBCS2* and *OsRBCS4* knockdown lines and no detectable RBCS protein in event *rbcs2*-266 and *rbcs4*-53 (Fig. 2B). There are coordinated reductions of RBCS and rbcL in both *rbcs2* and *rbcs4* lines and rbcL protein were reduced about 28% in *rbcs2*-165, 41% in *rbcs2*-266, 50% in *rbcs4*-022 and 39% *rbcs4*-53 compared to wild-type plants (Fig. 2A; Table 1). Immunolocalization confirmed reduction of Rubisco protein preferentially in the MCs (Fig. S3). Reduction in Rubisco content was correlated with the number of antisense inserts. *rbcs2*-266 and *rbcs4*-022 carried four and three antisense inserts, respectively, and their Rubisco protein contents were lower comparable to the events carrying fewer antisense inserts.

**Table 1.** Rubisco large subunit protein content.

|  | Percentage<br>abundance to WT |  |  |  |
| --- | --- | --- | --- | --- |
| *rbcs2*-165 | 72 | ± | 0.20 | * |
| *rbcs2*-266 | 59 | ± | 0.19 | * |
| *rbcs4*-022 | 50 | ± | 0.25 | * |
| *rbcs4*-053 | 61 | ± | 0.30 | * |

Values are the average ± SE of one leaf from four plants of *rbcs2* and *rbcs4* knockdown lines and wild-type (WT). Asterisk denotes statistically significant differences with WT, Student t-test, *p*-value ≤0.05.

**Fig. 1.**
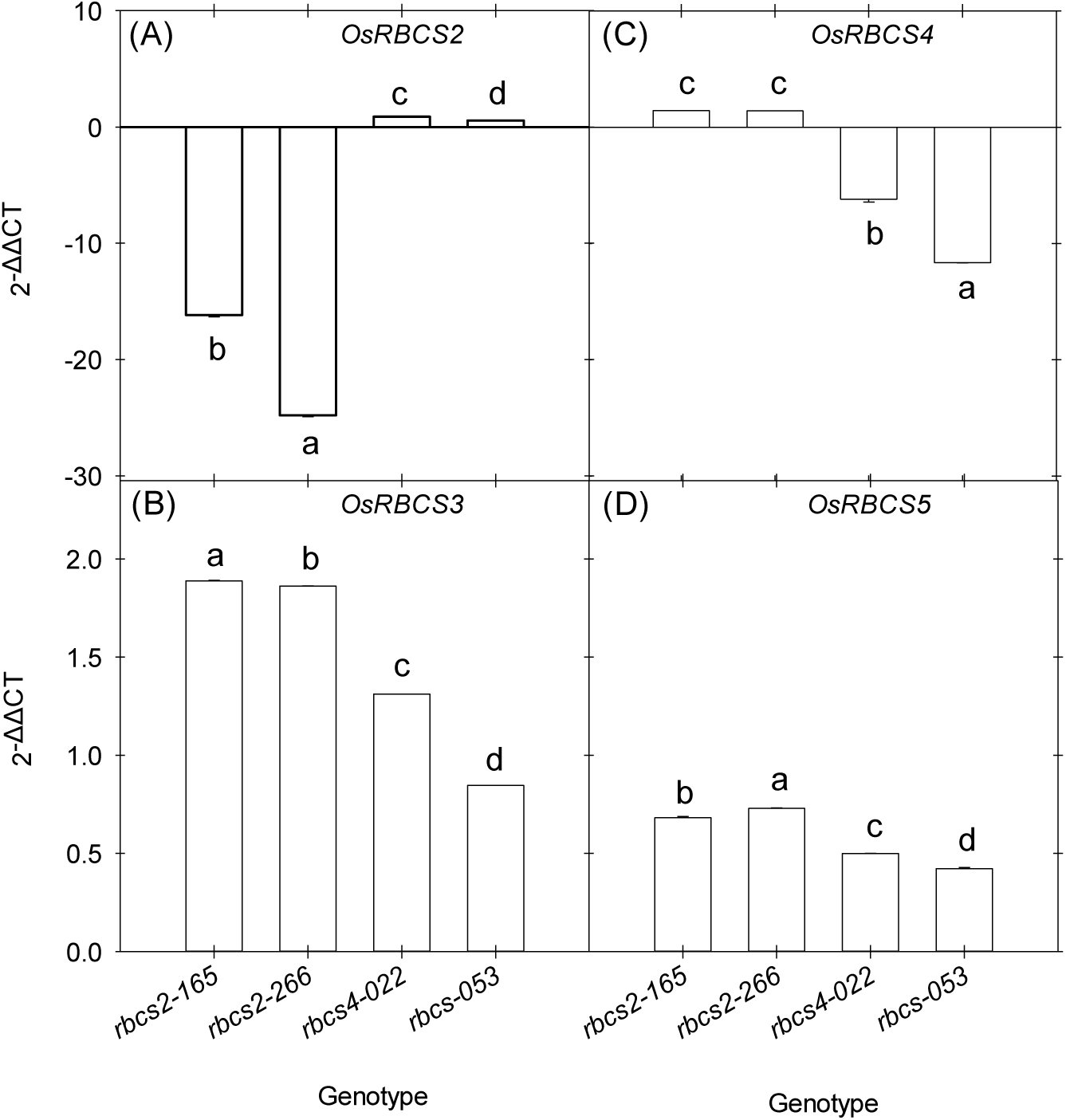
Transcript accumulation of four *OsRBCS* isoforms (*OsRBCS*2-5) as measured by quantitative real-time PCR in the leaf blade of *rbcs* knockdown lines. Values are expressed as fold changes in transcript accumulation relative to wild-type (mean ± SE). 2^−ΔΔCT^ was used to quantify relative abundance against *OsEF-1α* transcript (Os03g0177500). Values are the mean ± SE of three leaves from three plants per genotype. Different letters within groups denote statistically significant differences, ANOVA *p-*value ≤0.05.

**Fig. 2.**
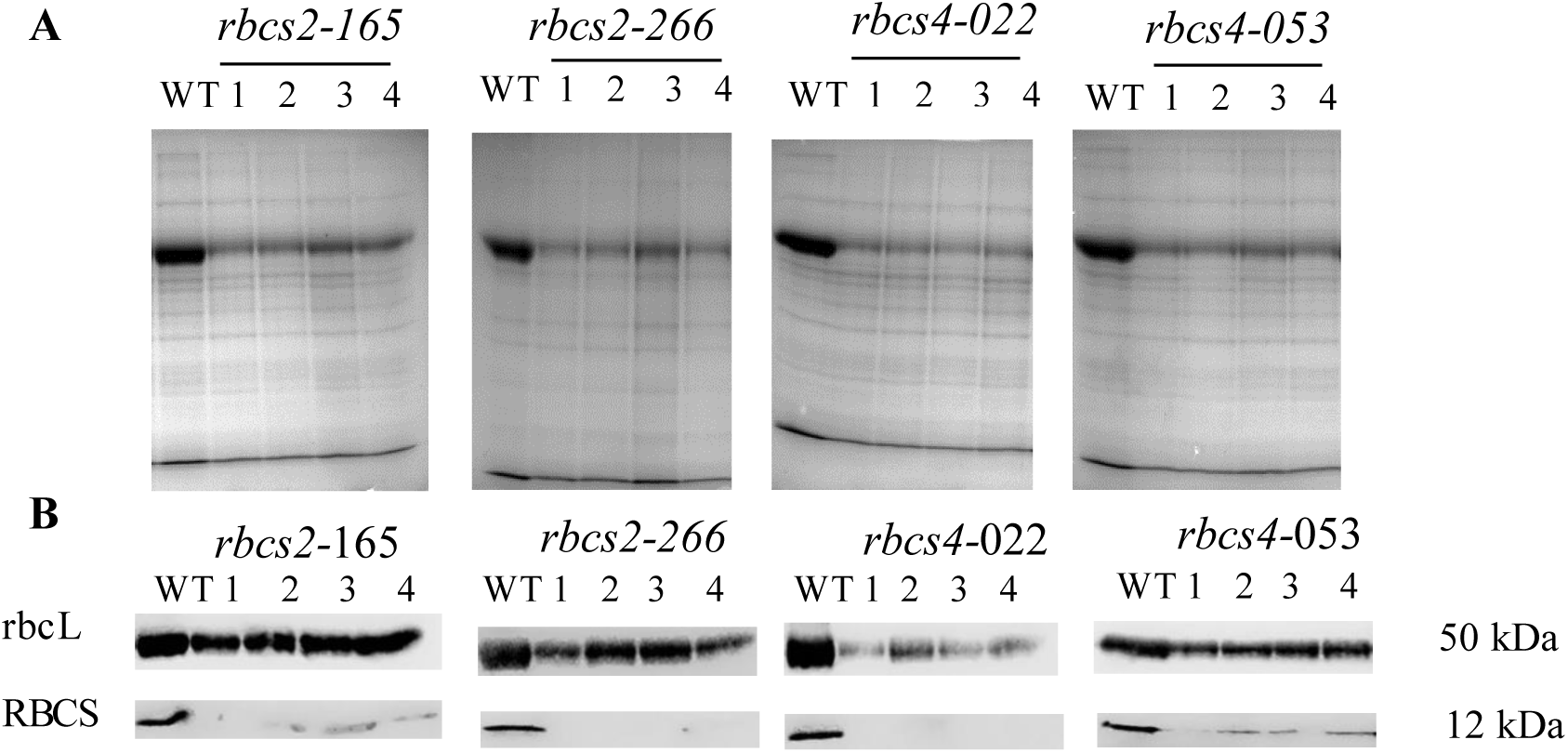
Soluble leaf protein. (A) SDS-PAGE and (B) immunoblots with antibodies against rbcL and RBCS. Wild-type (WT) and plants of *rbcs2* and *rbcs4* knockdown lines (numbers). Total extracted proteins were separated by 12% SDS-gel. Samples were loaded based on equal leaf area of 3.96 mm^2^.

### Photosynthetic perturbations associated with reduced Rubisco

The response of the net rate of CO_2_ assimilation (*A*) to intercellular CO_2_ concentration (*Ci*) was measured under photorespiratory (21% O_2_; Fig. 3A) and non-photorespiratory conditions (2% O_2_; Fig. 3B) at high irradiance (2,000 µmol photon m^−2^ s^−1^) where RuBP-saturated rates of Rubisco are limiting to photosynthesis. Under photorespiratory conditions, *A* was significantly lower in both *rbcs2* and *rbcs4* lines at all intercellular CO_2_ concentrations compared to wild-type rice, although photosynthesis was not saturated in any of the lines. Rates of CO_2_ assimilation were much lower in *rbcs4*-022 and *rbcs2*-226 lines compared to the other lines for the same gene; this was related to percentage reduction in Rubisco protein as *rbcs4*-22 and *rbcs2*-226 had lowest Rubisco content among four *rbcs* knockdown lines (Fig. 1). The responses under non-photorespiratory conditions were similar to those under photorespiratory conditions, except for a slight decrease in curvature and convergence in *rbcs2-*165 and wild-type response at the highest intercellular CO_2_ concentrations (Fig. 3B). These results suggest that photosynthesis was limited by Rubisco content under almost all conditions. Consistent with this, carboxylation efficiency (*CE*) was significantly reduced under both conditions, but most markedly under photorespiratory conditions (Table 2), indicative of Rubisco limited capacity in the leaf. CO_2_ compensation points (ᴦ) were significantly higher in event *rbcs2*-266 and *rbcs4*-022 compared with wild-type plants under both conditions (Table 2). In response to changes in irradiance, CO_2_ assimilation in wild-type plants was not saturated at 2,000 µmol photon m^−2^ s^−1^ (Fig. 3C). In the knockdown lines, photosynthesis was saturated at a of PPFD ̴1000 µmol photon m^−2^ s^−1^, with CO_2_ assimilation rates less than half those of wild-type plants. There was no discernable relationship between CO_2_ assimilation rate and Rubisco protein reduction, although there were line specific differences, with notably higher rates in *rbcs4-053* compared to other antisense lines. Quantum efficiency (Φ) and respiration rates (R_*d*_) were statistically significantly lower in all *rbcs* lines compared with wild-type plants (Table 2).

**Table 2.** Comparison of photosynthesis parameters.

| | $r$<br>$\mu\text{mol CO}_2 \text{ m}^{-2} \text{ s}^{-1}$ | $CE$<br>$\mu\text{mol CO}_2 \text{ m}^{-2} \text{ s}^{-1}$<br>$\mu\text{mol CO}_2 \text{ mol}^{-1}$ | $R_d$<br>$\mu\text{mol CO}_2 \text{ m}^{-2} \text{ s}^{-1}$ | $\Phi$ | |
| --- | --- | --- | --- | --- | --- |
|  |  |  |  |  | 21 % O<sub>2</sub> |
| Wild-type | $51.69 \pm 1.59^c$ | $0.126 \pm 0.02^a$ | $0.53 \pm 0.05^a$ | $0.046 \pm 0.003^a$ | |
| *rbcs2*-165 | $51.92 \pm 0.15^c$ | $0.067 \pm 0.0^b$ | $0.41 \pm 0.03^b$ | $0.039 \pm 0.002^b$ | |
| *rbcs2*-266 | $66.72 \pm 1.45^a$ | $0.046 \pm 0.0^b$ | $0.39 \pm 0.07^b$ | $0.031 \pm 0.000^b$ | |
| *rbcs4*-022 | $57.77 \pm 1.58^b$ | $0.05 \pm 0.01^b$ | $0.35 \pm 0.02^b$ | $0.030 \pm 0.000^b$ | |
| *rbcs4*-053 | $49.85 \pm 2.71^c$ | $0.06 \pm 0.01^b$ | $0.48 \pm 0.05^b$ | $0.025 \pm 0.000^b$ | |
|  |  |  |  |  | 2 % O<sub>2</sub> |
| Wild-type | $13.32 \pm 0.95^c$ | $0.166 \pm 0.04^a$ | - | - | |
| *rbcs2*-165 | $14.67 \pm 1.11^c$ | $0.086 \pm 0.0^b$ | - | - | |
| *rbcs2*-266 | $24.88 \pm 1.54^a$ | $0.057 \pm 0.0^b$ | - | - | |
| *rbcs4*-022 | $19.54 \pm 0.71^b$ | $0.06 \pm 0.01^b$ | - | - | |
| *rbcs4*-053 | $14.24 \pm 3.12^c$ | $0.08 \pm 0.01^b$ | - | - | |

CO<sub>2</sub> compensation point ( $r$ ), carboxylation efficiency ( $CE$ ), respiration rate ( $R_d$ ) and quantum efficiency ( $\Phi$ ) of wild type and *rbcs2* and *rbcs4* antisense lines. Measurements for $CE$ and $r$ were made at a photosynthetic photon flux density (PPFD) of 2,000 $\mu\text{mol photons m}^{-2} \text{ s}^{-1}$ , leaf temperature 30 °C and either 21% or 2% O<sub>2</sub>. Measurements of $R_d$ were made after 30 min in the dark. Measurements of $\Phi$ were made at a CO<sub>2</sub> concentration of 400 $\mu\text{mol CO}_2 \text{ mol}^{-1}$ and a leaf temperature of 30 °C. Values represent the mean ± SE of three leaves from three plants per genotype. Different letters within groups indicate statistical significance from a one-way ANOVA ( $p \leq 0.05$ ) and a least significant difference test (LSD) for post-hoc pairwise comparison.

**Fig. 3.**
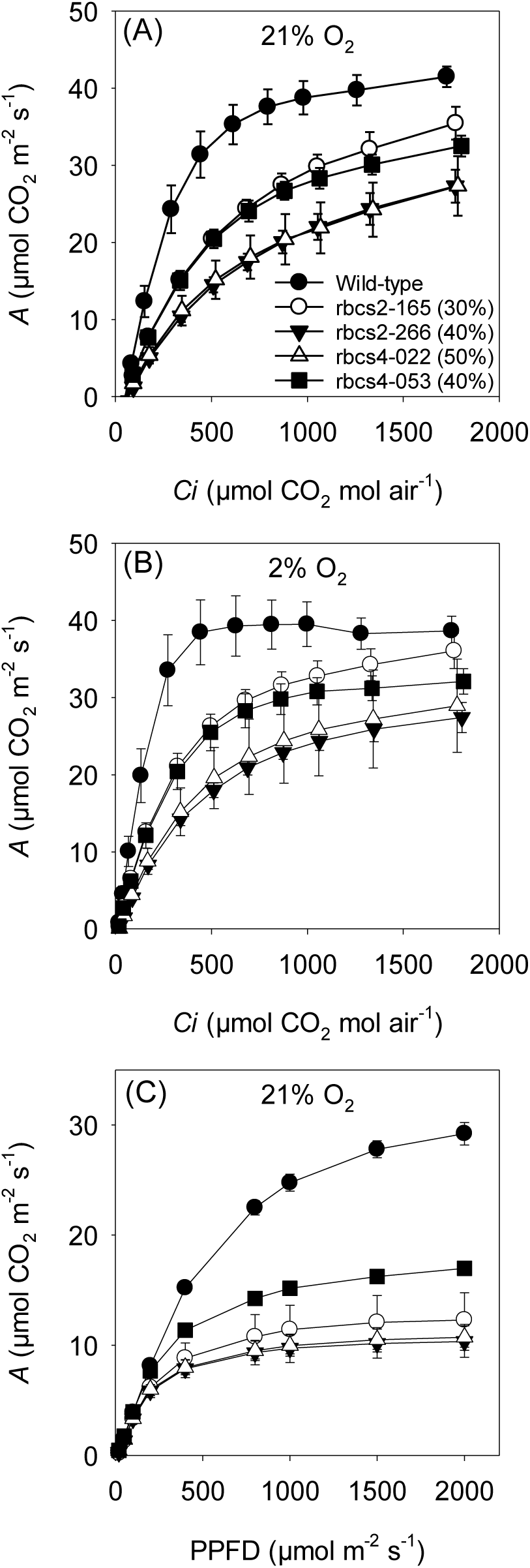
Net CO_2_ assimilation rate (*A*) in wild-type and *rbcs2* and *rbcs4* knockdown lines. In response to intercellular CO_2_ concentration (C*i*) measured at 21% O_2_ and (B) 2% O_2_. Measurements were made at a light intensity of 2,000 µmol photons m^−2^ s^−1^ and a leaf temperature of 30 °C. In response to photosynthetic photon flux density (PPFD; C). Measurements were made at CO_2_ concentration of 400 µmol CO_2_ mol air^−1^ and a leaf temperature of 30 °C. Values in legend are percentage reduction in Rubisco protein compared to the wild-type. Values are the mean ± SE of three leaves from three plants per genotype.

### Phenotypic perturbations associated with reduced Rubisco

Reduction of Rubisco protein in *rbcs2* and *rbcs4* knockdown lines led to a statistically significant reduction in grain yield of between 37 to 70% (Table 3), with considerable variation observed between individual lines. In all lines except *rbcs2-*165, there was a corresponding statistically significant reduction in dry biomass, with *rbcs4* lines exhibiting the largest decrease relative to the wild-type plants. There were no statistically significant differences in tiller number but *rbcs4* lines were slightly shorter than wild-type plants (Table 3). There were no consistent differences in leaf chlorophyll contents, with small but statistically significant reductions in *rbcs4* lines and a higher chlorophyll content in *rbcs2*-266. The difference in phenotype perturbations between lines follows protein abundance with *rbcs2*-165 exhibiting the lowest Rubisco protein reduction and smallest phenotypic perturbations and *rbcs4-*022 the largest Rubisco protein reduction and most significant phenotypic perturbations.

**Table 3.** Leaf chlorophyll content, plant height, tiller number, dry biomass and grain yield of wild-type and *rbcs* knockdown lines.

|  | Chl<br>(SPAD value) | Plant height<br>cm | Tiller number | Dry biomass<br>g | Grain yield<br>g |
| --- | --- | --- | --- | --- | --- |
| Wild-type | 43.14 ± 0.75<sup>b</sup> | 105 ± 1.4<sup>a</sup> | 26.1 ± 4.5<sup>abc</sup> | 58.16 ± 10.93<sup>a</sup> | 28.84 ± 7.68<sup>a</sup> |
| *rbcs2-165* | 42.30 ± 0.71<sup>bc</sup> | 105.6 ± 1.9<sup>a</sup> | 29.2 ± 3.9<sup>a</sup> | 57.36 ± 6.81<sup>a</sup> | 18.07 ± 5.59<sup>b</sup> |
| *rbcs2-266* | 44.04 ± 0.93<sup>a</sup> | 103.6 ± 1.2<sup>a</sup> | 27.5 ± 4.4<sup>ab</sup> | 44.32 ± 5.12<sup>b</sup> | 14.33 ± 4.52<sup>b</sup> |
| *rbcs4-022* | 41.84 ± 1.20<sup>c</sup> | 101.4 ± 1.8<sup>b</sup> | 22.6 ± 4.9<sup>bc</sup> | 32.60 ± 8.76<sup>c</sup> | 8.90 ± 2.44<sup>b</sup> |
| *rbcs4-053* | 42.00 ± 0.97<sup>c</sup> | 99.4 ± 3.0<sup>b</sup> | 23.4 ± 4.8<sup>c</sup> | 33.19 ± 8.23<sup>c</sup> | 13.80 ± 5.39<sup>b</sup> |

Chlorophyll contents (SPAD value) are the average ± SE of three leaves from ten plants per genotype. Plant height, tiller number, dry biomass and grain yield are the average ± SE of ten individual plants per line; 77 days post-germination. Biomass is the total dry weight of leaf, stem and sheath tissue. Different letters within groups indicate statistical significance based on a one-way ANOVA ( $p \leq 0.05$ ) with a Least Significant Difference (LSD) test for post-hoc pairwise comparison.

### Effect of reduced Rubisco on leaf anatomy

There were statistically significant differences in interveinal distance (IVD) between individual *rbcs* knockdown lines (Table 4), with *rbsc2*-165 having the widest IVD and *rbcs4*-022 line the lowest. Mesophyll cell number was significantly reduced in all the *rbcs* knockdown lines. The mesophyll cell length was increased in all *rbcs* knockdown lines but was only statistically significant in *rbcs2*-165 (Table 4). No difference in the number and thickness of major or minor veins were found. Leaves of both *rbcs4* knockdown lines were significantly narrower than either the wild-type or *rbcs2* lines, which correspond with reduced IVD and MC number.

**Table 4.**
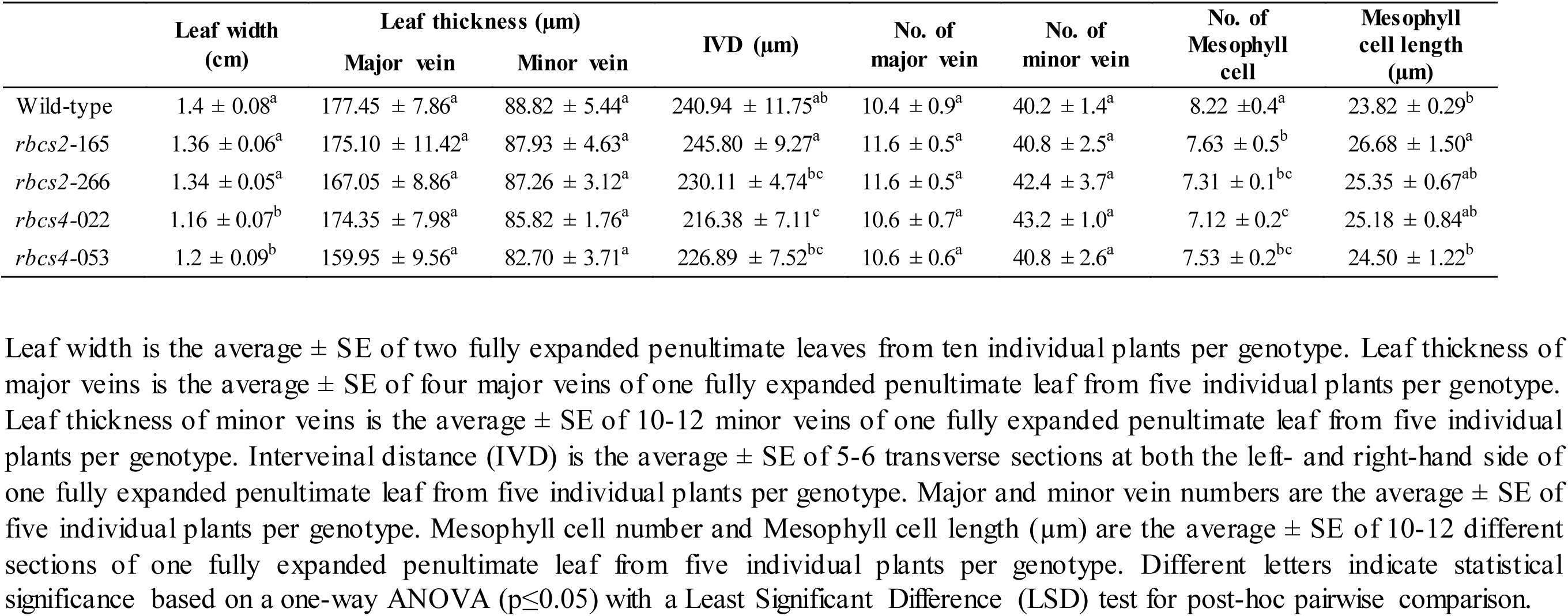
Leaf width, leaf thickness, interveinal distance (IVD), vein number and mesophyll cell characteristics of wild-type and *rbcs* knockdown lines.

|  | Leaf width<br>(cm) | Leaf thickness (µm) |  | IVD (µm) | No. of<br>major vein | No. of<br>minor vein | No. of<br>Mesophyll<br>cell | Mesophyll<br>cell length<br>(µm) |
| --- | --- | --- | --- | --- | --- | --- | --- | --- |
|  |  | Major vein | Minor vein |  |  |  |  |  |
| Wild-type | 1.4 ± 0.08<sup>a</sup> | 177.45 ± 7.86<sup>a</sup> | 88.82 ± 5.44<sup>a</sup> | 240.94 ± 11.75<sup>ab</sup> | 10.4 ± 0.9<sup>a</sup> | 40.2 ± 1.4<sup>a</sup> | 8.22 ± 0.4<sup>a</sup> | 23.82 ± 0.29<sup>b</sup> |
| *rbcs2-165* | 1.36 ± 0.06<sup>a</sup> | 175.10 ± 11.42<sup>a</sup> | 87.93 ± 4.63<sup>a</sup> | 245.80 ± 9.27<sup>a</sup> | 11.6 ± 0.5<sup>a</sup> | 40.8 ± 2.5<sup>a</sup> | 7.63 ± 0.5<sup>b</sup> | 26.68 ± 1.50<sup>a</sup> |
| *rbcs2-266* | 1.34 ± 0.05<sup>a</sup> | 167.05 ± 8.86<sup>a</sup> | 87.26 ± 3.12<sup>a</sup> | 230.11 ± 4.74<sup>bc</sup> | 11.6 ± 0.5<sup>a</sup> | 42.4 ± 3.7<sup>a</sup> | 7.31 ± 0.1<sup>bc</sup> | 25.35 ± 0.67<sup>ab</sup> |
| *rbcs4-022* | 1.16 ± 0.07<sup>b</sup> | 174.35 ± 7.98<sup>a</sup> | 85.82 ± 1.76<sup>a</sup> | 216.38 ± 7.11<sup>c</sup> | 10.6 ± 0.7<sup>a</sup> | 43.2 ± 1.0<sup>a</sup> | 7.12 ± 0.2<sup>c</sup> | 25.18 ± 0.84<sup>ab</sup> |
| *rbcs4-053* | 1.2 ± 0.09<sup>b</sup> | 159.95 ± 9.56<sup>a</sup> | 82.70 ± 3.71<sup>a</sup> | 226.89 ± 7.52<sup>bc</sup> | 10.6 ± 0.6<sup>a</sup> | 40.8 ± 2.6<sup>a</sup> | 7.53 ± 0.2<sup>bc</sup> | 24.50 ± 1.22<sup>b</sup> |

Leaf width is the average ± SE of two fully expanded penultimate leaves from ten individual plants per genotype. Leaf thickness of major veins is the average ± SE of four major veins of one fully expanded penultimate leaf from five individual plants per genotype. Leaf thickness of minor veins is the average ± SE of 10-12 minor veins of one fully expanded penultimate leaf from five individual plants per genotype. Interveinal distance (IVD) is the average ± SE of 5-6 transverse sections at both the left- and right-hand side of one fully expanded penultimate leaf from five individual plants per genotype. Major and minor vein numbers are the average ± SE of five individual plants per genotype. Mesophyll cell number and Mesophyll cell length (µm) are the average ± SE of 10-12 different sections of one fully expanded penultimate leaf from five individual plants per genotype. Different letters indicate statistical significance based on a one-way ANOVA ( $p \leq 0.05$ ) with a Least Significant Difference (LSD) test for post-hoc pairwise comparison.

## Discussion

To mimic the downregulation of part of the Calvin–Benson cycle in MCs that is required for C_4_ photosynthesis, we have successfully targeted two *OsRBCS* genes, *OsRBCS2* and *OsRBCS4*. Genetic redundancy within the *OsRBCS* gene family does not completely compensate for a reduction in the expression of a single multigene family member (Kanno et al., 2017, Ogawa et al., 2012). Constructs designed to independently reduce expression of *OsRBCS2* and *OsRBCS4* in the MCs led to a significant reduction in gene transcript for the target gene with modest increases in the other multigene family members (Fig. 1). These results are consistent with previous studies showing that the expressions of each *OsRBCS* genes were regulated independently from other *OsRBCS* genes (Kanno et al., 2017, Ogawa et al., 2012). In leaf blades of wild-type plants *OsRBCS2* transcripts accumulate to similar levels as those of *OsRBCS4* (Suzuki et al., 2009). However, the effect of suppression of *OsRBCS2* antisense was stronger with a larger reduction in *OsRBCS2* transcript accumulation in *rbcs2* knockdown lines. In *rbcs2* knockdown lines Rubsico protein accumulation was 30-40% lower, which is consistent with reports from previous studies where Rubisco content was reduced by up to 45% in *OsRBCS2* cDNA antisense rice lines (Makino et al., 1997; Makino et al., 2000). Although we did not generate lines targeting all the *OsRBCS* multigene family, these results suggest that targeting multiple genes is required to completely reduce Rubisco accumulation in MCs.

Reducing Rubsico protein content by 30% or more in the antisense rice plants limits CO_2_ assimilation rates under photorespiratory conditions (21% O_2_). A relationship between the reduction in Rubisco protein content and CO_2_ assimilation rate was observed, which is consistent with previous studies (Makino et al., 1997; Suzuki et al., 2012). On the other hand, other studies also reported that a relatively small decrease in Rubisco content (65-90% of wild-type Rubisco) can lead to an increase in CO_2_ assimilation rate of between 5–15% at elevated CO_2_ due to the excessive amount of Rubisco for photosynthesis in rice (Kanno et al., 2017; Makino et al., 1997; Makino et al., 2000). However here both *rbcs2* and *rbcs4* knockdown lines did not have higher CO_2_ assimilation rate than wild-type plants in elevated CO_2_ conditions. The differences between results in this study and previous studies may due to reduction of *OsRBCS2* and *OsRBCS4* being preferentially targeted to the MCs leading to a more severe reduction in Rubisco content in the MCs. On a protein quantity basis, the reduction of *rbcs2* lines had a more significant impact on CO_2_ assimilation rate than *rbcs4* lines given the percentage reduction in Rubisco protein.

A reduction in Rubisco protein content and CO_2_ assimilation was associated with a significant reduction in dry biomass accumulation and grain yield. The magnitude of the reduction was proportional to the reduction in Rubisco protein content. The *rbcs* knockdown lines had no difference in number of major and minor veins or leaf thickness compared to wild-type plants (Table 4). Evan et al. (1994) reported a small reduction (92% of wild-type) in leaf thickness in the *rbcs* antisense tobacco lines when Rubisco was removed by more than 50%. We also observed that the numbers of MCs in between minor veins was reduced and the mesophyll cell length was slightly increased in *rbcs* knockdown lines (Table 4). This suggests that there may be anatomical changes to the leaf associated with altering Rubisco protein content.

Our results show that preferentially MC targeted antisense-*rbcs* plants can be used to reduce Rubisco accumulation in rice to levels that induce a phenotype without lethality. The resulting knockdown plants have reduced CO_2_ assimilation rates with moderate effects on plant growth. This provides a useful foundation for installing C_4_ photosynthesis in rice.

## Abbreviations

*A*: net rate of CO_2_ assimilation
*ACi*: photosynthesis CO_2_ response curves
BSC: bundle sheath cell
*C*_*a*_: ambient CO_2_ concentration
CE: carboxylation efficiency
Chl: chlorophyll
*C*_*i*_: intercellular CO_2_ concentration
rbcL: large subunit of Rubisco
MC: mesophyll cell
PPFD: photosynthetic photon flux density
R_d_: dark respiration rate
Rubisco: ribulose-1,5-bisphosphate carboxylase/oxygenase
RuBP: ribulose 1,5-bisphosphate
RBCS: small subunit of Rubisco

---

**Fig. S1.**
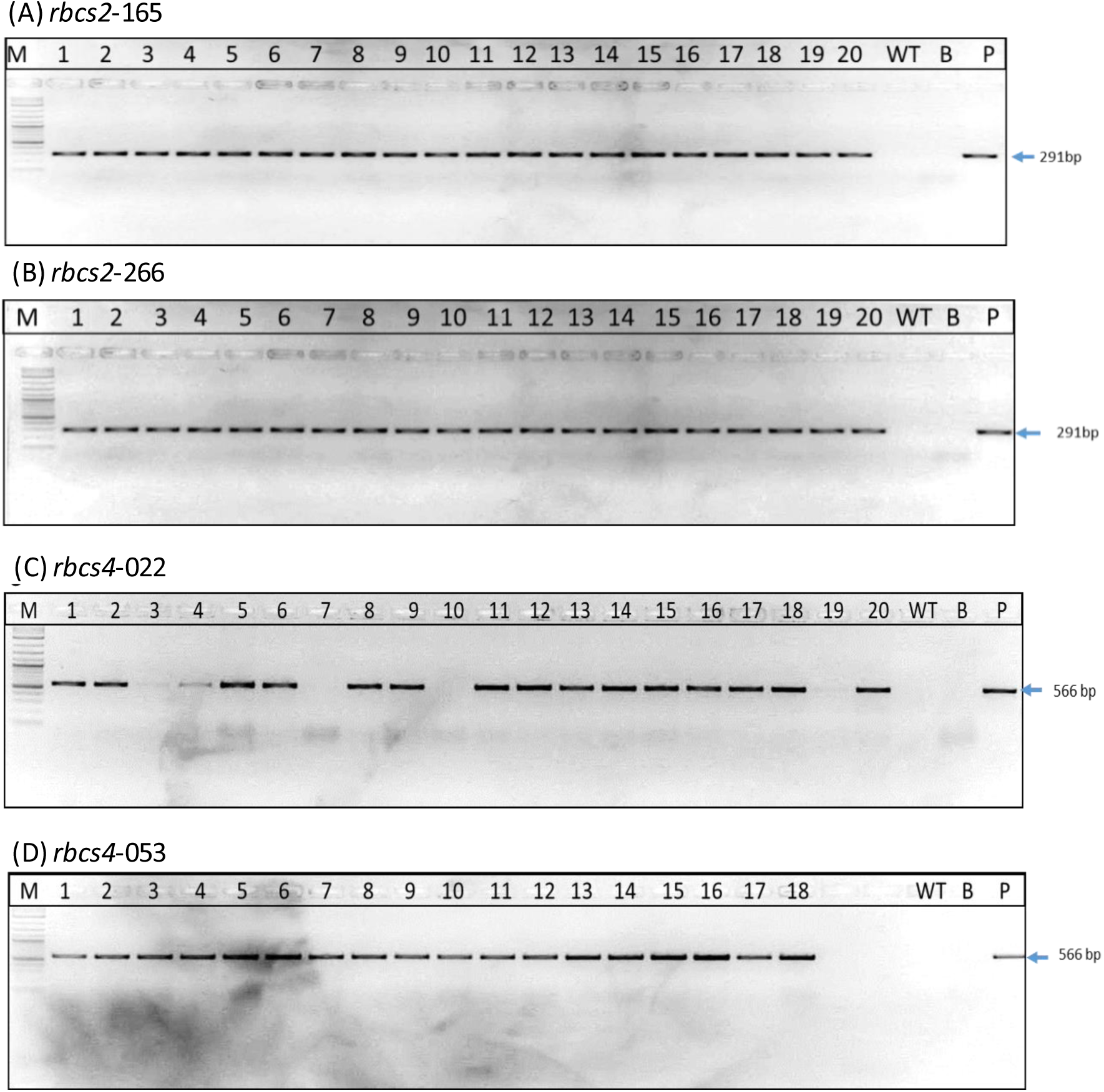
PCR analysis of *rbcs2* and *rbcs4* knockdown lines (numbers). Wild-type (WT) was used as a negative control and the vector plasmid (P) as positive control. B indicates blank (H_2_O), M indicates a ladder (100bp).

**Fig. S2.**
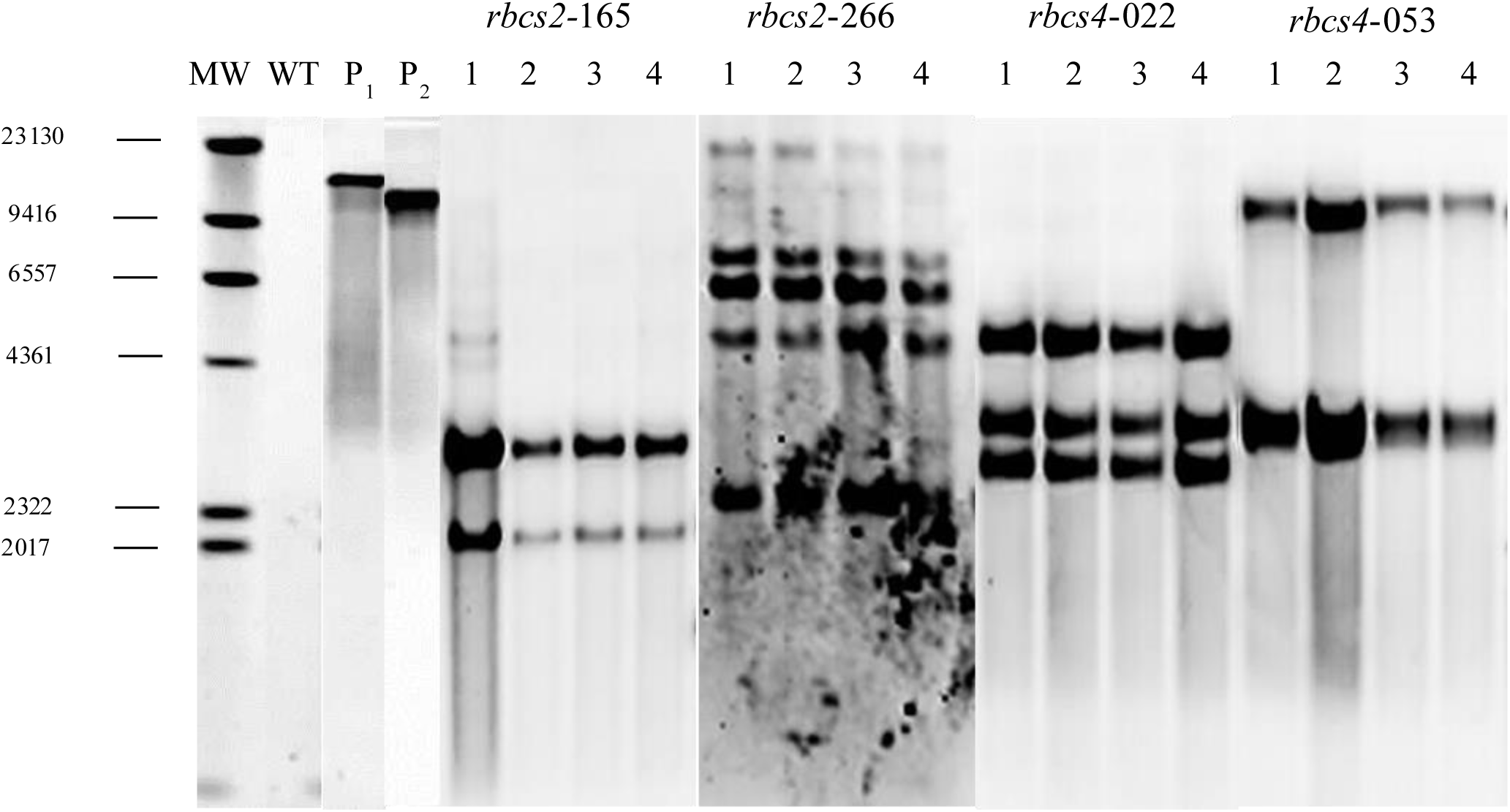
DNA blot analysis of *rbcs2* and *rbcs4* knockdown lines (numbers). The genomic DNA of each plant was extracted from leaves with the CTAB method and digested with *BgL*II. Wild-type plants (WT) were used as a negative control and the vector plasmid (P1 for antisense-*OsRBCS2* construct and P2 for antisense*-OsRBCS2* construct) as a positive control. A DIG-labeled DNA molecular marker indicates the molecular weight (MW).

**Fig. S3.**
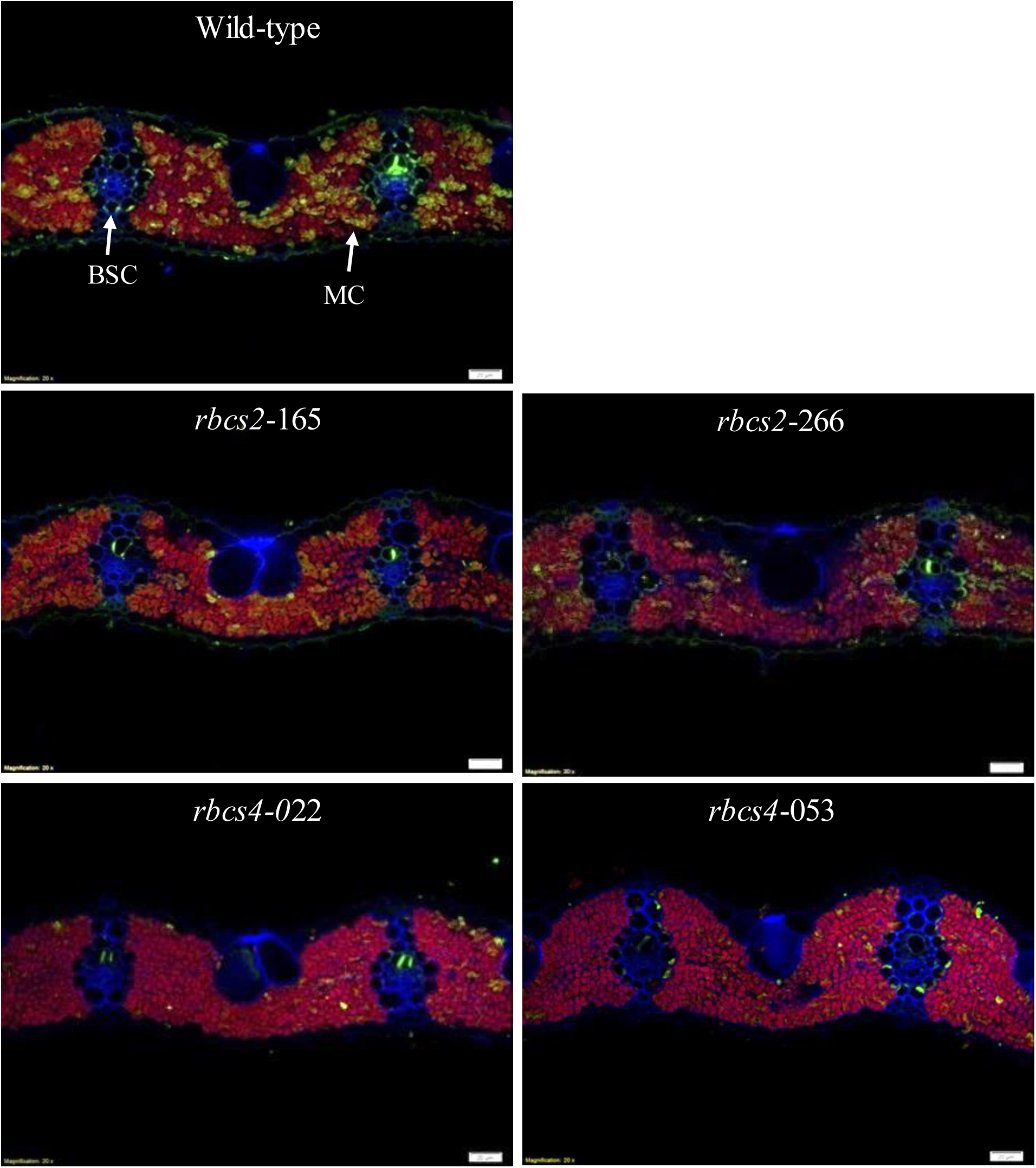
Representative images of immunolocalization of Rubisco protein in leaves from wild-type (WT), *rbcs2* and *rbcs4* knockdown lines. Anti-Rubisco rabbit polyclonal primary antibody diluted 1:200 plus Alexa Fluor 488 goat anti-rabbit IgG as secondary antibody diluted 1:200 was used to probe for rbcL (shown in green color). Chlorophyll is seen as a red autofluorescence. The cell wall was visualized by co-staining with calcofluor white and is shown in blue. Magnification: 200x. Scale bar: 20 μm. BSC: Bundle sheath cell. MC: mesophyll cell.

## Notes

### Competing Interest Statement

The authors have declared no competing interest.

